# DIMT1, a regulator of ribosomal biogenesis, controls β-cell protein synthesis, mitochondrial function and insulin secretion

**DOI:** 10.1101/2020.09.27.313262

**Authors:** Gaurav Verma, Alexander Bowen, Alexander Hamilton, Jonathan Esguerra, Alexandros Karagiannopoulos, Luis Rodrigo Cataldo, Claire Lyons, Elaine Cowan, Malin Fex, Hindrik Mulder

## Abstract

We previously reported that transcription factor B1 mitochondrial (TFB1M) is involved in the pathogenesis of type 2 diabetes (T2D) owing to mitochondrial dysfunction. Here, we describe that dimethyladenosine transferase 1 homolog (DIMT1), a homologue of TFB1M, is expressed and active in pancreatic β-cells. Like TFB1M, it has been implicated in control of ribosomal RNA (rRNA) but its role in β-cells or T2D remains to be identified. Silencing of *DIMT1* impacted mitochondrial function, leading to reduced expression of mitochondrial OXPHOS proteins, reduced oxygen consumption rate (OCR), dissipated mitochondrial membrane potential (ΔΨm) and caused a lower rate of ATP production (mATP). In addition, *DIMT1* knockdown slowed the rate of protein synthesis. In accordance with these findings, DIMT1-deficiency perturbed insulin secretion in rodent and human β-cell lines. These effects are likely a result of destabilization of ribosomal assembly, involving NIN1 (RPN12) binding protein 1 homolog (NOB-1) and Pescadillo ribosomal biogenesis factor 1 (PES-1). These are two critical ribosomal subunits proteins, whose interactions were perturbed upon DIMT1-deficiency, thereby disturbing protein synthesis in β-cells. Thus, we have here highlighted a role of DIMT1 in ribosomal biogenesis that perturbs protein synthesis, resulting in mitochondrial dysfunction and disrupted insulin secretion, both being potential pathogenetic factors in T2D.

## Introduction

Type 2 Diabetes (T2D) is the result of dual defects of insulin secretion and insulin action (Basu *et al*, 2009; Cavaghan *et al*, 2000). Pancreatic β-cells compensate for insulin resistance by secreting increased amounts of insulin. Over time, in predisposed individuals, β-cells become exhausted and fail to maintain adequate insulin levels to overcome insulin resistance and maintain blood glucose. The rising hyperglycemia, insulin resistance and hyperinsulinemia may also in themselves contribute to metabolic abnormalities (Lee *et al*, 2017; Shanik *et al*, 2008). Insulin secretion from the β-cell is controlled by glucose metabolism (Schuit *et al*, 1997; Wollheim & Maechler, 2002). Following a meal, glucose is taken up by transporters, before being metabolized via glycolysis and the citric acid cycle. In these events, the mitochondrion is a critical player, accounting for a major part of cellular metabolism (Fex *et al*, 2018). Importantly, mitochondrial metabolism produces ATP, which closes ATP-sensitive K^+^ channels, leading to depolarization of the plasma membrane (Ashcroft, 2005). Subsequently, voltage-dependent Ca^2+^ channels open, allowing Ca^2+^ entry, which ultimately triggers insulin secretion (Gandasi *et al*, 2017; Rorsman *et al*, 2012). In addition, mitochondria produce a number of potential coupling factors, which amplify secretion of insulin (Brun & Maechler, 2016; Fex *et al*, 2018). Therefore, β-cell function and insulin secretion are critically dependent on mtDNA and protein expression. (Koeck *et al*, 2011; Nicholas *et al*, 2017).

We have reported that a variant of the gene encoding transcription factor B1 mitochondrial (TFB1M) is associated with reduced insulin secretion, hyperglycemia and future risk of T2D (Koeck *et al.*, 2011). TFB1M was initially considered to act as a transcription factor along with TFB2M but more recent reports support a role as methyltransferase (Cotney & Shadel, 2006; Seidel-Rogol *et al*, 2003) (Metodiev *et al*, 2009). In mitochondria, TFB1M dimethylates two adjacent adenine residues in 12S rRNA, conferring stability to the mitochondrial ribosome (Guja *et al*, 2013). Our studies showed that TFB1M-deficiency leads to mitochondrial dysfunction and impaired insulin secretion resulting in diabetes (Sharoyko *et al*, 2014). Despite this important role of RNA regulation and structure, the role of other methyltransferases in RNA methylation has not been investigated in β-cells. Therefore, we set out to examine functions of other methyltransferases that could be implicated in β-cell function and T2D.

To this end, we identified DIMT1 (Dimethyladenosine Transferase 1 Homolog), which shows 50% homology with TFB1M, and is expressed in human islets and clonal β-cells. DIMT1 is a well-known methyltransferase involved in ribosome biogenesis but its function in β-cells has not been explored. Using rodent and human insulin-producing cell lines, we show that DIMT1 regulates protein synthesis, impacting on mitochondrial function and consequently insulin secretion. Thus, we have identified a possible role of DIMT1 in β-cell ribosome biogenesis and mitochondrial function.

## Material and Methods

### Cell lines and reagents

EndoC-βH1 cells (Endo Cells, Paris, France) were grown on Matrigel-fibronectin coated (100 μg/mL and 2 μg/mL, respectively, Sigma-Aldrich, Steinheim, Germany) cell culture plates in DMEM containing 5.6 mM glucose, 2% BSA fraction V (Roche Diagnostics, Mannheim, Germany), 10 mM nicotinamide (Merck Millipore, Darmstadt, Germany), 50 μM 2-mercaptoethanol, 5.5 μg/mL transferrin, 6.7 ng/mL sodium selenite (Sigma-Aldrich), 100 U/mL penicillin, and 100 μg/mL streptomycin (PAA Laboratories, Pasching, Austria). INS-1 832/13 cells were cultured as described (Hohmeier *et al*, 2000). The protein estimation kit used was from Pierce, Thermo Scientific, (Rockford, IL). All primary antibodies were from Abcam (Cambridge, United Kingdom) and the appropriate secondary antibodies used were from Santa Cruz (Santa Cruz Biotechnology, Inc., Dallas, Texas, USA). All the reagents used for qRT-PCR were from Thermo Scientific, (Rockford, IL). The TaqMan quantitative real-time PCR (qRT-PCR) master mix was from Applied Biosystems, CA, USA. The control (scramble) and human *DIMT1* siRNA (Cat# 4392420 and 4392421) were from Thermo Scientific, (Rockford, IL) and Rat *DIMT1* siRNA (Cat# NM_001106408 (SASI_Rn02_00218668) was from (Sigma Aldrich, St. Louis, Missouri, United States) Fluorescent probes were supplied by Molecular Probes (Rockford, IL). Insulin was measured by an ELISA kit (Mercodia, Uppsala, Sweden). All other chemicals were from Sigma Aldrich.

### Cell Incubations

EndoC βH1-cells were grown in DMEM culture medium with 5.6 mM glucose. INS-1 832/13 cells were cultured in RPMI medium (11.1 mM glucose). Both cell lines were incubated in the absence and presence of of *DIMT1* siRNA at 0, 50, 100 and 200 nM concentrations for 48 or 72h; cells were transfected by Lipofectamine RNAi Max (Invitrogen, CA, USA) according to the manufacturer’s instructions. We used pre-designed DIMT1 siRNA (Thermo Scientific) Cat# 4392420 and 4392421 for EndoC-βH1 cells (Sense:GGAUGGUCUAGUAAGGAUAtt), (Antisense:UAUCCUUACUAGACCAUCCca) and Cat# SASI_Rn02_00218668 (Sigma Aldrich) for INS-1 832/13 cells (Sense:CUGUUCAGUACAGAAUACUdt), (Antisense:AGUAUUCUGUACUGAACAGdt). For NBR1 we used Cat# SASI_Hs01_00169473, (Sense:GCUUAAGAUGGCAGUUAAA[dT][dT]) and (Antisense: UUUAACUGCCAUCUUAAGC[dT][dT]) and for DNAJC19 Cat# SASI_Hs01_00055864, (Sense:CAGCAUUAAUACUAGGUGU[dT][dT]) and (Antisense:ACACCUAGUAUUAAUGCUG[dT][dT]) (Sigma Aldrich). On termination of incubation, cells were assessed for cell viability with trypan blue; other assays were performed as described below.

### RNA isolation and Quantitative Real-Time PCR

Total RNA was extracted from the cells, using TRI Reagent (Sigma Aldrich) according to manufacturer’s instructions. RNA concentrations were determined by NanoDrop Spectrophotometer (Thermo Scientific). Equal quantities of total RNA were reversely transcribed using RevertAid First-Strand cDNA synthesis kit (Fermentas, Vilnius, Lithuania) in reactions containing 500 ng of total RNA. qRT-PCR was performed using TaqMan gene expression assays (Human-Assay ID Hs00917510_m1; and (Rat-Assay ID Rn01489483_m1; Applied Biosystems, Life Technologies, Carlsbad, CA). Gene expression was quantified by the comparative Ct value, in which the amount of target is expressed as 2^−ΔΔCt^ using actin as a reference gene.

### Western blot analysis

Cells were washed with phosphate-buffered saline (PBS) and homogenized in cell lysis buffer (150mM NaCl, 1% NP40, 10% DOC, 10% SDS, 50mM Tris, pH 7.4). Determination of protein concentration was performed using the BCA protein assay kit (Pierce, Rockford, IL). Samples were mixed with standard Laemmli loading buffer and 30 μg protein loaded on mini-Protean TGX precast gel (Bio-Rad, USA). After SDS-PAGE, proteins were transferred to Immunoblot PVDF membranes (BioRad), the membrane blocked with 3% BSA, and further incubated with rabbit monoclonal anti-DIMT1 antibody (1:1000 dilution) and Rabbit polyclonal anti-OXPHOS (1:250 dilution antibody; both from Abcam, Cambridge, UK). Tubulin antibody (1:5000 dilution; Abcam, Cambridge, UK) was used as a loading control. Horseradish peroxidase-linked goat anti-rabbit IgG (1:5000 dilution; Santa-Cruz Biotechnology) was used as secondary antibody. Blots were developed with enhanced chemiluminescence (ECL). Densitometry analysis was performed using BioRad ImageLab software.

### Insulin Secretion and Content

Confluent EndoC-βH1 cells were starved in medium at 2.8 mM glucose overnight and incubated next day in 1X secretion assay buffer (SAB) (114 mM NaCl, 1.2 mM KH_2_PO_4_, 4.7 mM KCl, 1.16 mM MgSO_4_, 25 mM CaCl2, 100 mM HEPES,25mM NaHCO_3_ 0.2% BSA, pH 7.4) containing 1 mM glucose for 2 h. Finally, cells were incubated in SAB containing the indicated glucose concentrations for 1 hour. After the incubation, supernatants and lysates were collected. Insulin secretion and content were measured by Human (#10-1113-01) and Rat Insulin ELISA Kit (#10-1250-01) Mercodia, Uppsala, Sweden) according to manufacturer’s instructions.

### Protein synthesis measurement

Human β-cells were seeded in a 96 well dish with a density of 5 × 10^4^ cells and transfected with *DIMT1* siRNA. At confluence, cells were harvested by centrifugation (400 × g, 5 min), and supernatant carefully removed; cells were resuspended in 500 μl medium containing 25 μM OPP (#601100, Cayman Chemicals, MI, USA). Subsequently, cells were incubated for 30 min at 37 °C and 5% CO_2_. After incubation, cells were centrifuged at (400 × g, 5 min) and supernatants were removed. One hundred μl of cell-based fixatives were added to each well and incubated for 5 mins at room temperature. Cells were then centrifuged and 100 μl of wash buffer was added, and cells were incubated for another 5 mins and centrifuged again. One hundred μl of 5 FAM-Azide solution was added and cells were incubated for 30 mins. After the final wash, cells were examined in fluorescent plate reader using a filter designed to detect FITC (excitation/emission= 485/535 nm). Three specificity controls were included: 1) cells labelled with OPP; 2) knock down cells labelled with OPP and; 3) cells incubated with cyclohexamide at a final concentration of 50 μg/mL for 15 min prior to, as well as during incubation with OPP.

### Primer extension assay to measure methylation

For the primer extension assay, sense and anti-sense primers were designed flanking the methylated adenosine residues on human 18s rRNA. Hemo KlenTaq polymerase was used, which specifically transcribes DNA but not the methylated DNA with secondary structures (Aschenbrenner & Marx, 2016). qRT-PCRs were performed in a total volume of 20 μl reaction mixture containing 100 nM of Taqman gene expression assay (Human-Assay ID Hs00917510_m1), 200 μM dNTPs (each), in KlenTaq reaction buffer. The concentrations of the different RNAs were quantified and 500 ng of each sample were reverse transcribed as explained before. The primer sets used for the detection of the methylated adenosine residues were: 1) *DIMT1* Forward *GACGGTCGAACTTGACTATCTA*; 2) *DIMT1* Reverse *AATGATCCTTCCGCAGGT*; and 3) probe *AGTCGTAACAAGGTTTCCGTAGGTGA*. Gene expression was quantified by the comparative Ct value, in which the amount of target is expressed as 2^−ΔΔCt^ using actin as a reference gene.

### Mitochondrial membrane potential (ΔΨm) measurement

Human β-cells were seeded onto collagen coated 8-well chambered cover glasses (Lab-Tek, Thermo Scientific) at a density of 70,000 cells/cm^2^. After 24 h, cells were transfected with 100 nM of *DIMT1* siRNA and incubated further for 72 h. For the ΔΨm measurement, cells were pre-incubated with imaging buffer (135 mM NaCl, 3.6 mM KCl, 1.5 mM CaCl_2_, 0.5 mM MgSO_4_, 0.5 mM Na_2_HPO_4_, 10 mM HEPES, 5 mM NaHCO_3_, pH 7.4) containing 2 mM glucose for 2 hr with 100 nM of tetramethylrhodamine methyl ester (TMRM; Invitrogen, CA, USA). After the incubation, cells were washed again with the imaging buffer only and subjected to live cell confocal microscopy in quench mode, where the whole cell fluorescence decreases upon mitochondrial hyperpolarization. After recording the basal level of TMRM fluorescence in 2 mM glucose, cells were switched to 20 mM glucose. Carbonyl cyanide-4-phenylhydrazone (FCCP) was used to dissipate the ΔΨm. Zeiss LSM510 inverted confocal fluorescence microscope with 543 nm excitation and 585 nm long pass emission settings were used to record the data, which were background corrected and normalized.

### ATP:ADP Measurement

Single cell ATP/ADP ratio measurements were carried out, using co-expression of a pericam-based ATP biosensor (Perceval HR; Addgene ID: 49083) and pHRed. EndoC-βH1 cells were seeded onto collagen coated 8-well chambered cover glasses (Lab-Tek, Thermo Scientific) at a density of 70,000 cells/cm^2^. After 24 h, cells were co-transfected with 1 μg of plasmid encoding Perceval HR (Addgene ID:49083) and 100 nM of *DIMT1 siRNA* at 50% cell confluency. Cells were grown for 72 h before measurements; cells were pre-incubated at 37°C in 400 μl of imaging buffer (135 mM NaCl, 3.6 mM KCl, 1.5 mM CaCl_2_, 0.5 mM MgSO_4_, 0.5 mM Na_2_HPO_4_, 10 mM HEPES, 5 mM NaHCO_3_, pH 7.4) containing 2 mM glucose. After recording the basal signal from Perceval HR (2 mM glucose), 20 mM of glucose was added. Cells were imaged with 490 nm excitation and 535 nm emission filter settings for Perceval and pHRed was excited using 578/16 and 445/20 nm band pass filters; the emissions were collected through a 629/56 nm band pass filter on Zeiss LSM510 inverted confocal fluorescence microscope.

### Respiration

Mitochondrial oxygen consumption rate (OCR) was determined in EndoC-βH1 cells, using the Seahorse Extracellular Flux Analyzer XF24 (Seahorse Bioscience, Billerica, MA). After transfection (48 h) with *DIMT1* or scramble siRNA, cell culture medium was exchanged for 500 μl of Seahorse assay buffer supplemented with 1 mM glucose for 2h at 37°C. OCR was recorded in intact cells stimulated by 10 mM pyruvate. Oligomycin (OM)-independent OCR (5 μg/ml) was measured. The mitochondrial inner membrane ionophore (FCCP, 4 μM) was added to determine maximal respiratory capacity and rotenone+Antimycin (1 μM) was added to block the transfer of electrons from complex I to ubiquinone. Data were analysed by the Seahorse wave software (Agilent, Santa Clara, CA 95051, US)

### Immunoprecipitation for binding partners

EndoC-βH1 cells were treated with *DIMT1* siRNA and lysates were isolated using a proprietary lysis buffer (Pierce, Thermo Scientific, (Rockford, IL). Co-IP was performed at 4°C unless otherwise indicated, using a Pierce spin column. The binding of NOB1 antibody to protein A/G agarose beads was performed according to the manufacturer’s instructions. Protein A/G agarose slurry (20 μl) was washed twice with 200 μl PBS buffer, and incubated with 100 μl NOB1 antibody prepared in PBS (10 μl NOB1 antibody +85 μl H_2_O+5 μl 20X PBS) at 25°C for 30 min on a gentle shaker. The supernatants were discarded and the beads washed three times with 300 μl PBS. After removing the supernatant, beads were washed three times again. The antibody-cross-linked beads were incubated overnight at 4°C with 500 μl of 1 mg lysates which were pre-cleared with control agarose resin (Pierce) for 2 h on a shaker. After removing supernatant (flow-through) and washing with 600 μl washing buffer five times, the immunoprecipitates were eluted with 40 μl 2XLaemmli buffer at 100°C for 10 min. Equal amounts of eluted input and co-IP lysates complex were subjected to SDS-PAGE separation for western blotting.

### Proximity ligation assay for protein-protein interaction

For proximity ligation assay (PLA), DuoLink PLA technology probes and reagents (Cat# DUO92101-1KT) (Sigma-Aldrich) were used. Cells were permeabilized using the combination of paraformaldehyde, methanol and ethanol incubation for 10 min. After two washes in PBS, cells were incubated with blocking solution for 30 min at 37°C and then with the two different primary antibodies (NOB1 and Pescadillo) for 1 h at room temperature. The cover slips were washed twice for 5 min with buffer A, followed by incubation with the PLA probes (secondary antibodies against two different species bound to two oligonucleotides: anti-mouse MINUS and anti-rabbit PLUS) in antibody diluent for 60 min at 37°C. After two washes of 5 min with buffer A, the ligation step was performed for 30 min at 37°C. The cells were washed with buffer A twice for 2 min before incubation for 100 min with amplification stock solution at 37°C. The amplification stock solution contains polymerase for the rolling circle amplification step and oligonucleotides labelled with fluorophores, which will bind to the product of the rolling circle amplification and thus allowing detection. After two washes of 10 min with buffer B, cells were incubated with FITC-conjugated buffers. Finally, the cover slips were washed with PBS and mounted with Duolink in situ mounting medium containing DAPI. The cells were visualized under the florescent microscopy with UV lasers for nucleus and TRITC for the detection of red PLA signals (excitation wavelength of 594nm and an emission wavelength of 624 nm).

### RNA Sequencing

RNA was extracted from EndoC-βH1 cells and sequencing was performed using Illumina RiboMethSeq Protocol (ebook ISBN 978-1-4939-6807-7, chapter 12). The library was prepared by NEB-Next multiplex small RNA library prep set for Illumina. The libraries were loaded and sequenced on Illumina NextSeq500 sequencer in the sequencing core facility at Lund University Diabetes Centre, Malmö, Sweden. Transcript reads were mapped to the human transcriptome (Gencode Release 30) and quantified with Salmon (v.0.14.0). Differentially expressed genes were identified with DESeq2 (v. 1.25.10).

### Panther pathway Analysis

Significantly altered genes were functionally classified via molecular function and biological processes, using the PANTHER classification system (http://www.pantherdb.org). The candidate gene list was converted into text file (ID list) with *Homo sapiens* selected as the organism database; functional classification was viewed in pie chart. PANTHER classifies genes on the basis of published experimental evidence to predict functions. To assess our dataset relative to the global set of human genes, binomial statistics and Fischer’s exact test for multiple testing within the PANTHER system was applied. An outline of the detailed study design is depicted in Supplementary (Figure 1).

**Figure 1:**
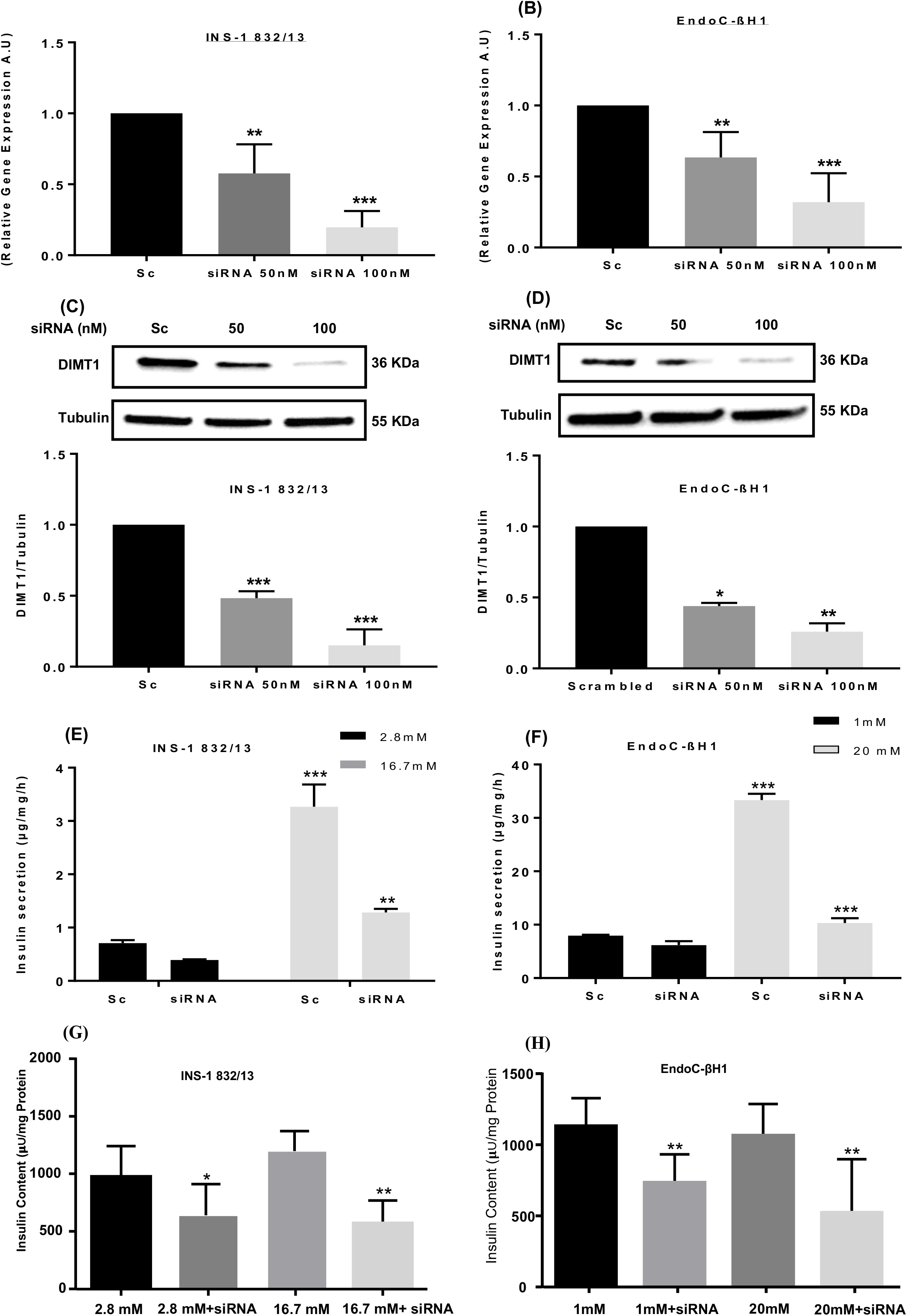
*DIMT1*-deficiency and its impact on insulin secretion. Gene expression of *DIMT1* is shown in silenced INS-1 832/13 (A) and EndoC-βH1 cells (B); cells treated with with scramble siRNA served as control and actin mRNA expression as internal control. *DIMT1* silencing at the protein level was determined by western blot in INS-1 832/13 (C) and EndoC-βH1 cells (D); tubulin was used as loading control. Data are expressed as mean ±S.E.M for both experiments (n=5, EndoC-βH1 and n = 5, INS-1 832/13). Insulin secretion at 2.8 mM glucose (black bar) and 16.7 mM glucose (grey bar) in INS-1 832/13 cells is shown in (E); insulin secretion at l 1mM (black bar) and 20 mM glucose (Grey bar) in EndoC-βH1cells (F). Insulin content is shown in both the cell types with upon glucose stimulation (G, H); total cellular protein was used for normalization both for insulin secretion and content. Data are expressed as mean ±S.E.M (n=3, INS-1 832/13 and n= 3, EndoC-βH1). Data were statistically compared with the paired Student’s t-test; *p<0.05, **p<0.01, ***p<0.001.

### Statistical analysis

Mean ± SE were calculated. Pair-wise comparisons were made by *chi-square*-test; to determine the statistical significance between two groups, parametric or non-parametric tests were used as indicated. A significance level of p < 0.05 was considered statistically significant.

## Results

### Knock down of *DIMT1* in INS-1 832/13 and EndoC-βH1 cells

We used siRNA duplexes targeting separate regions of *DIMT1* mRNA in both INS-1 832/13 and EndoC-βH1cells. (Figure 1A-B) shows knock down of *DIMT1* mRNA in the cell lines post-transfection; (Figure 1C-D) shows knock down of DIMT1 protein. One hundred nM siRNA effectively silenced mRNA and protein levels by ~80% both in INS-1 832/13 and EndoC-βH1 cells 72 hr post-transfection. We, therefore, used 100 nM of siRNA in all of our subsequent experiments.

### *DIMT1* knock down impairs insulin secretion in β-cells

To understand whether loss of DIMT1 influences insulin secretion and content, we investigated the effect of siRNA-mediated *DIMT1* knock down on glucose-stimulated insulin secretion (GSIS) and insulin content in INS-1 832/13 and EndoC-βH1cells. Stimulation with 16.7 mM glucose led to a 3-fold increase in insulin secretion in INS-1 832/13 cells; in EndoC-βH1 cells, 20 mM glucose led to a 4-fold increase in insulin release as compared to low glucose (Figure 1E-F). In contrast, *DIMT1* knock down in both cell lines resulted in nearly a 3-fold decrease in insulin release in response to high glucose compared to control (Figure 1E-F). No significant differences were observed in insulin content between low and high glucose concentrations in either cell line, however, *DIMT1* knock down significantly reduced the insulin content in both INS-1 832/13 and EndoC-βH1 cells when compared with cells treated with scrambled siRNA (Figure 1G-H).

### DIMT1 methylates 18S rRNA in β-cells

*DIMT1* has been reported to be responsible for N^6^,N^6^ adenosine dimethylation at positions A_1850_ and A_1851_ on human 18S RNA (Zorbas *et al*, 2015). To examine whether *DIMT1* also serves as a dimethylase in human β-cells, we performed primer extension assays with primers flanking the methylation region on human 18S rRNA. As shown in (Figure 2A), Hemo KlenTaq failed to transcribe mRNA isolated from untransfected EndoC-βH1 cells or cells treated with scrambled siRNA. In contrast, knock down of *DIMT1* (50nM and 100nM) concentration-dependently reinstated the expression of 18S rRNA due to abrogation of DIMT1 methylation activity. We conclude that *DIMT1* specifically methylates the 18S rRNA in EndoC-βH1 cells (Figure 2A).

**Figure 2:**
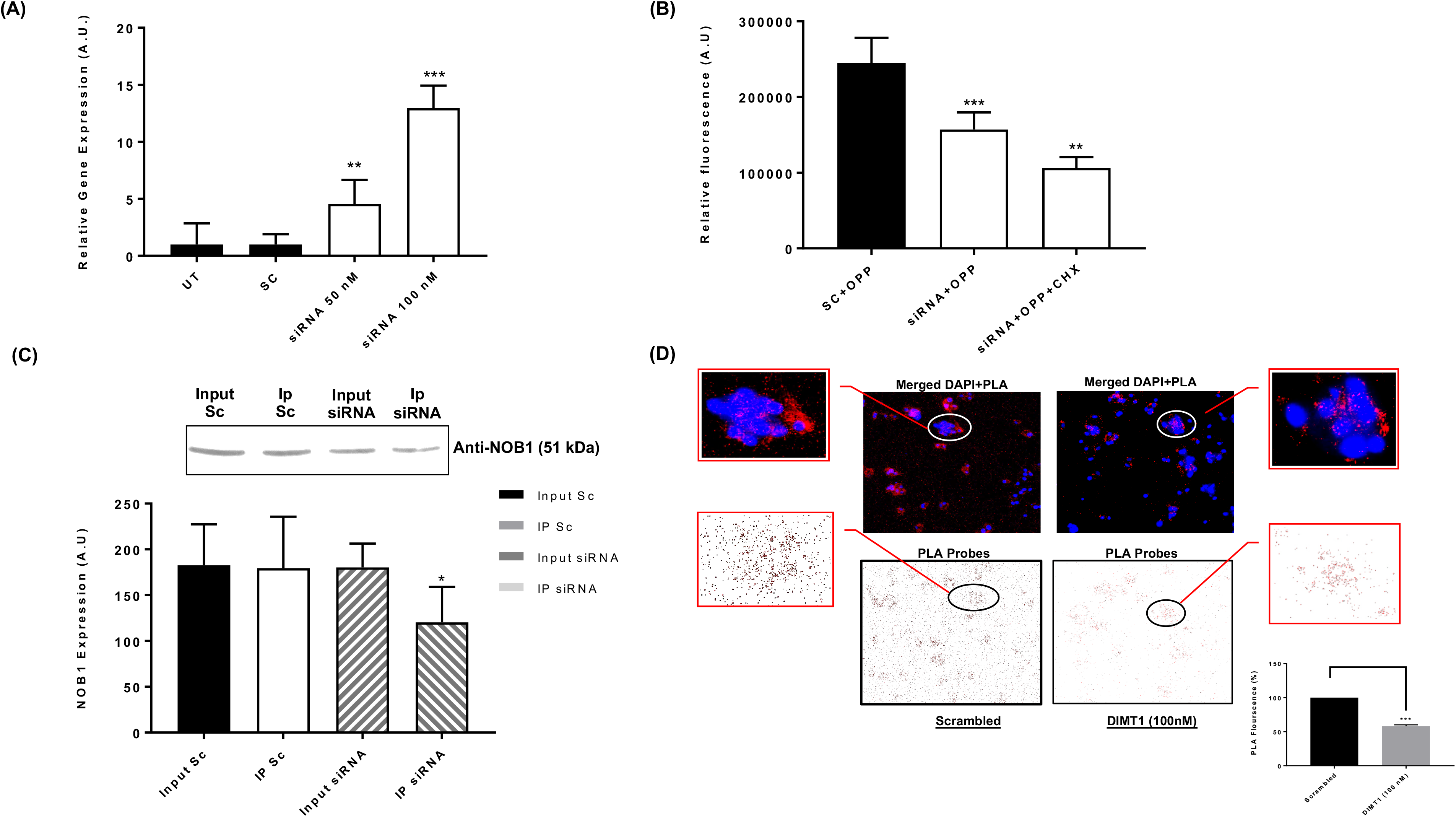
The role of DIMT1 role in 18S methylation and protein expression. Quantification of 18S ribosomal methylation from total RNA was made by use of qRT-PCR (A). The black and white bar indicates control cells and cells treated with *DIMT1* siRNA, respectively. Data are expressed as mean ±S.E.M (n = 4). Quantification of protein synthesis with OPP fluorescence in (B); black bar shows the Alexa568 fluorescence intensity in control and the white bar shows the reduced OPP fluorescence in *DIMT1* siRNA- and cycloheximide (CHX)-treated cells. Data are expressed as mean ±S.E.M (n = 3). Physical interaction of NOB1 and PES-1 is shown in (C) with immunoprecipitation expressed as mean ±S.E.M (n = 3). (D) Proximity ligation assay (PLA) for NOB-1/PES-1 interaction is measured by red dot signals that represent the proximity of the NOB-1 and PES-1 proteins. DNA is stained with DAPI. Scale bar: 10 μM. Quantification of the PLA signals per cell after *DIMT1* siRNA treatment is shown in the graph where, the data are expressed as mean ±S.E.M (n = 3); statistical comparisons between *DIMT1*-silenced cells and scramble control cells were made by Student’s t-test; **p<0.01, ***p<0.001.

### DIMT1-deficiency perturbs protein synthesis in β-cells

Ribosomes are required for protein synthesis in cells. To examine whether reduced 18S rRNA methylation impacts protein synthesis, we assayed protein synthesis in *DIMT1*-silenced EndoC-βH1 cells with a puromycin-based assay. Addition of puromycin labelling led to a drastic reduction of mRNA translation rates under CHX treatment (Figure 2B). As a consequence, we observed that puromycin addition led to less total protein synthesis upon *DIMT1* knock down, as compared to the scramble control. Cells treated with the scrambled siRNA and puromycin (SC+OPP) showed a significant increase in OPP fluorescence, reflecting increased global protein synthesis rates in these cells (Figure 2B),. In contrast, DIMT1-deficient cells showed a decreased protein synthesis rate (Figure 2B). Of note, overall protein synthesis was examined, including mitochondrial protein synthesis. At this point, it was not possible to distinguish between cellular or mitochondrial protein synthesis rates. Nevertheless, we conclude that DIMT1 plays a role in β-cell protein synthesis.

### DIMT1 perturbs assembly of ribosomal subunits in EndoC-βH1 cells

So far, we observed that DIMT1-deficiency impairs protein synthesis, insulin secretion and content, but the underlying molecular mechanism remains to be determined. Recent reports have suggested that DIMT1 and WBSCR22 methylate the 18S rRNA, which is required for pre-rRNA processing reactions leading to stabilization of the ribosomal subunit (Zorbas *et al.*, 2015). Therefore, we further investigated the role of DIMT1 in ribosomal biogenesis and protein synthesis. We used immunoprecipitation (IP) and a proximity ligation assay (PLA) to test whether DIMT1-deficiency perturbs protein translation in EndoC-βH1 cells by destabilizing the ribosomal subunit. To this end, we chose two critical rRNA proteins from each subunit of the 40S and 60S rRNA for IP: NOB-1 (Nin1,binding protein), an endonuclease involved in the assembly of 40S rRNA, primarily required for cleavage of the 20S pre-rRNA to generate the mature 18S rRNA, and PES-1 (pescadillo ribosomal biogenesis factor 1), an essential ribosomal biogenesis factor of the 60S subunit. As shown in (Figure 2C), PES-1 was able to pull down NOB-1 protein from the EndoC-βH1 cell lysates both in scrambled controls and in *DIMT1*-silenced cells. Interestingly, we observed decreased levels of NOB-1 in DIMT1-deficient samples (Figure 2C, lane 4). This suggests that DIMT1 is playing a crucial role in ribosomal assembly because NOB-1 is an integral part of a well-defined pre-40S ribosomal complex containing the 20S precursor to the 18S rRNA (Woolls *et.al.* 2011).

To strengthen and substantiate our findings in EndoC-βH1 cells, we performed a PLA. We probed the control and *DIMT1*-silenced cells with Duolink florescent PLA probes, and used two different primary antibodies to probe for NOB-1 and PES-1. We observed an increased number of PLA probes predominantly in the cytoplasm (the number of each probe represents the number of the interactions) in scramble controls as compared with *DIMT1*-deficient EndoC-βH1cells (Figure 2D). We did not detect any signal from the nucleus, visualized by DAPI-staining, suggesting that the ribosomal maturation process is largely cytoplasmic. We found lower levels of probes in *DIMT1*-silenced cells as compared to the scramble siRNA-treated cells (45% in *DIMT1* knock down cells). This indicates that the two proteins associated strongly in the presence of DIMT1 whereas their interactions were diminished in DIMT1-deficient cells. This circumstance is likely to interfere with protein translation. Collectively, based on our observations, DIMT1-deficiency appears to destabilize the 40S subunit, which then fails to interact with 60S, leading to reduced protein translation rate. The latter notion is supported by our finding of reduced protein synthesis in the puromycin-based assay and reduced insulin content.

### DIMT1-deficiency perturbs mitochondrial function in β-cells

Given that *DIMT1* knock down perturbed insulin secretion stimulated by glucose in EndoC-βH1 cells, it is likely that DIMT1-deficiency disrupts mitochondrial control of stimulus-secretion coupling in β-cells. To this end, we first investigated the mitochondrial oxidative phosphorylation complexes (OXPHOS), using a panel of selected antibodies to both mitochondrial and nuclear-encoded proteins. We found decreased levels of OXPHOS protein expression in *DIMT1*-silenced EndoC-βH1 cells (Figure 3A). Levels of two subunits of respiratory complexes, complex V (ATP synthase) and the complex III mitochondrial-encoded (Cyt b), were markedly reduced in *DIMT1*-silenced cells (Figure 3A). In addition, DIMT1-deficiency also led to a significant decrease of protein levels in the nuclear encoded subunits of complex II (succinate dehydrogenase) and complex IV, (COX1). These results suggest that DIMT1 may play a crucial role in protein biosynthesis of mitochondrial OXPHOS proteins, which could contribute to mitochondrial dysfunction. These data also suggest that mitochondrial ribosomal biogenesis was impacted by DIMT1-deficiency in EndoC-βH1 cells. The graphical representation of the OXPHOS blots is shown in (Figure 3B).

**Figure 3:**
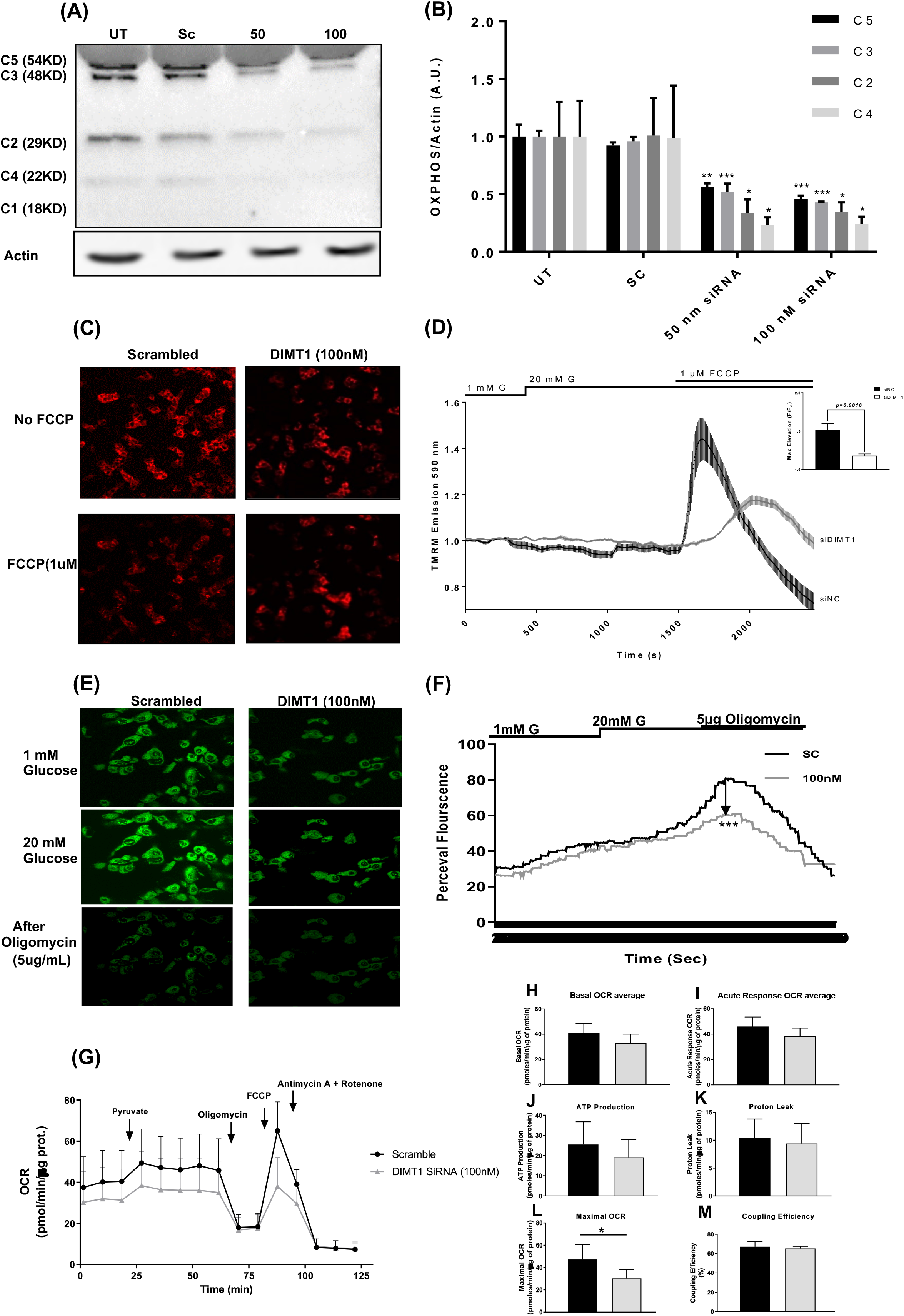
DIMT1 and Mitochondrial Dysfunction in EndoC-βH1 cells. Representative immunoblots (A) and densitometric analyses (B) of mitochondrial complex protein I-V. Data are means ± SEM of values from three independent experiments. Glucose-induced (1mM and 20mM) mitochondrial hyperpolarization of the inner mitochondrial membrane are shown by TMRM (C); average traces (thick lines) in scramble cells compared to the difference in maximal hyperpolarization with FCCP in *DIMT1*-silenced cells (thin lines) are shown (D). Data (mean ± SEM) are from three independent experiments. Cytosolic ATP/ADP ratios determined by Perceval HR (green fluorescence) are shown in (E); average traces and the difference in maximal rise in ATP/ADP ratio with Oligomycin are shown in (F). Data (mean ± SEM) are from three independent experiments. (G) Mitochondrial OCR is shown upon stimulation with pyruvate (10 mM; G) as scramble cells (black lines) and as *DIMT1*-silenced cells (grey lines). Pyruvate-stimulated respiratory response, proton leak, ATP production and maximal mitochondrial respiration with FCCP and non-mitochondrial respiration (Antimycin+Rotenone) are each expressed as fold relative to basal (H-M); data are mean ±S.E.M (n= 4). Statistical differences were compared by Student’s t-test. *p<0.03 or < 0.05 (as indicated), **p < 0.01 and ***p < 0.001.

Respiratory proteins control the electron transport chain, leading to proton extrusion over the inner mitochondrial membrane. Next, we examined the effect of *DIMT1* knock down on the inner mitochondrial membrane potential (ΔΨm), using a fluorescent dye (tetramethyl-rhodamine methylester; TMRM), which is sequestered by active mitochondria and reflects the ΔΨm. We observed, upon stimulation with 20 mM glucose, a decrease in TMRM fluorescence intensity from baseline in *DIMT1*-deficient cells as compared to scrambled controls (Figure 3C and 3D). In cells treated with the mitochondrial uncoupler FCCP, an abrogation of the increase in TMRM fluorescence intensity from baseline in scrambled control was evident when compared to *DIMT1*-silenced cells (Figure 3C and 3D). This suggests that *DIMT1* knock down dissipated the ΔΨm in EndoC-βH1 cells.

As *DIMT1* knock down led to decreased expression of ATP synthase (Figure 3A), we asked whether *DIMT1* knock down influenced the levels of mitochondrial ATP production in response to glucose. Using the ATP:ADP sensor Perceval HR, we observed a blunted increase in the ATP/ADP ratio, reflecting the rate of mitochondrial ATP production, in DIMT1-deficient cells (Figure 3E and 3F). In contrast, however, the scrambled control cells showed a robust increase in the ratio of ATP/ADP following a rise in glucose from 1 to 20 mM (Figure 3E). Addition of 5 μg/ml oligomycin, an ATP synthase blocker, reduced the ATP/ADP ratio regardless of treatment. Changes in pH may bias the Perceval HR signal; therefore, data were normalized by the pHRed traces. Our data suggest that disruption of DIMT1 in β-cells leads to altered expression of OXPHOS complexes leading to impaired mitochondrial ATP production.

Next, we used pyruvate as a substrate for mitochondrial metabolism, thereby avoiding effects exerted by glycolysis. Cells treated with scramble siRNA showed a significant rise in OCR in response to 10 mM pyruvate (Fig. 3G); respiratory complex inhibitors (oligomycin and antimycin + rotenone) reduces the OCR whereas the mitochondrial membrane uncoupler FCCP increases it in response to high metabolic control of ATP synthesis on OCR (highly coupled) and high maximal mitochondrial capacity respectively (Fig. 3G, J, L and M). While the *DIMT1*-deficient cells showed a similar pattern of OCR in response to pyruvate, maximal OCR was significantly reduced as compared to control cells (Fig. 3G; p=0.03; n = 3). This result is in line with the defects of OXPHOS proteins by *DIMT1*-silencing. Collectively, our results confirm that *DIMT1*-silenced cells exhibit perturbed mitochondrial function that culminates in impaired metabolism and reduced insulin secretion in EndoC-βH1 cells.

### DIMT1 targets in β-cells revealed by RNA sequencing

To gain insights into the effects of *DIMT1*-deficiency in β-cell transcriptome and to identify possible targets for this protein, we performed RNA sequencing in EndoC-βH1 cells. The sequencing data revealed 48 differentially expressed genes, out of which six genes whose expression was upregulated and six downregulated were selected for validation of our RNA sequencing results. As shown in (Figure 4A), mRNA expression of all the six downregulated genes was lower when analysed by RT-PCR in *DIMT1*-silenced cells. Similarly, we validated upregulation of five of the six selected upregulated genes identified by RNA sequencing (Figure 4B). *CTSH* (Cathepsin H) expression was found to be nominally upregulated but this did not reach statistical significance. Differentially expressed downregulated and upregulated genes analysis are shown in a volcano plot (Figure 4C).

**Figure 4:**
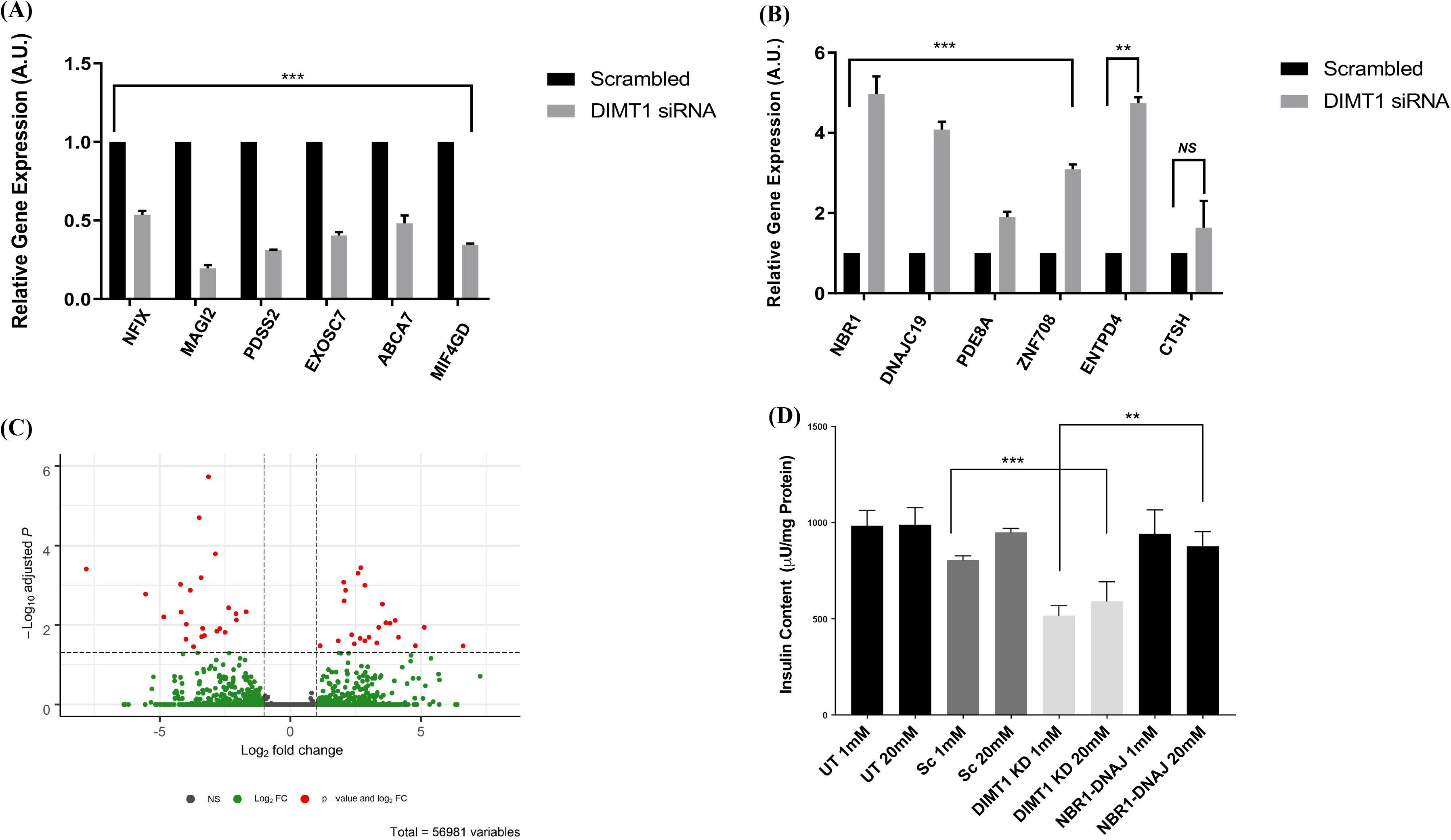
Validation of RNA sequencing by quantitative real-time PCR. Differentially expressed genes identified by RNA sequencing were evaluated; a total of six each differentially expressed genes were selected on the basis of their (i) fold change, (ii) relevance to cellular pathways and (iii) expression levels shown in (A and B). Data are mean ±S.E.M (n=5). (C) Volcano plot representing significantly downregulated genes (log2fold change< (−1) and adjusted p-value < 0.05), which are indicated in red to the left side of the plot, while significantly upregulated genes (log2fold change > (1) and adjusted p-value < 0.05) are indicated in red to the right of the plot. Vertical dashed lines correspond to the log_2_ fold change threshold of |1| and horizontal dashed line corresponds to the adjusted p-value threshold of 0.05 represented as–log_10_ adjusted p-value. (D) Changes in insulin content in *NBR1-* and *DNAJC19*-silenced cells for the validation of 28S and 5S were confirmed (E and F). Graphical representation shown of the qRT-PCR analysis and the data are expressed as mean ±S.E.M (n = 3). Statistical analysis was done using paired Student’s t-test. **p<0.01, ***p<0.001, NS – non-significant.

Differentially expressed genes were categorised by PANTHER analysis with respect to molecular functions and biological processes that includes catalytic activity, transcription/translation regulatory activity, binding function, biological regulation, biogenesis and metabolic process (Supplementary Figure 1). Of note, most differentially expressed genes were involved in mitochondrial function and cellular metabolism. *DNAJC19* is a mitochondrial interacting protein and plays a key role in mitochondrial function whereas *NAUK1* (NUAK family SNF1-like kinase 1) is required for Ca^2+^-dependent AMPK activity. Other upregulated genes, like *ZNF708* (a zinc finger protein transcription factor) and *ENTPD4* (Ectonucleoside Triphosphate Diphosphohydrolase 4) were also involved in metabolic pathways.

Interestingly, we found that two of the upregulated genes, *NBR1* (neighbor of BRCA1 gene 1) and *DNAJC19* (DnaJ (Hsp40) homolog, subfamily C, member 19), are also found in the human Mitocarta inventory of genes (Human Mitocarta 2.0, Broad Institute), suggesting a possible role of DIMT1 in mitochondrial function. Downregulated genes were also reported to be critically involved in various mitochondrial functions, protein synthesis and cellular metabolism pathways.

Since, we observed that *DIMT1* knock down reduced insulin content (Figure 1G-H), we reasoned that knock down of upregulated genes secondary to DIMT1-deficiency, as identified by RNA-Sequencing, could rescue the observed reduction in insulin content in EndoC-βH1 cells. As shown in (Figure 4D), knock down of the two upregulated genes *NBR1* and *DNAJC19* significantly hindered the reduction in insulin content observed in cells upon *DIMT1* knock down. This result suggests that knock down of *DIMT1* upregulates *NBR1* and *DNAJC19* and that this could interfere with insulin biosynthesis and hence content, a situation prevented by silencing of *NBR1* and *DNAJC1*.

It has previously been reported that *DIMT1* methylation of the 18S rRNA is required for normal protein translation in HEK-293 cells (Zorbas *et al.*, 2015). We show here for the first time the role of *DIMT1* in β-cells translation in EndoC-βH1 cells. We next used our unbiased and global RNA sequencing data to interrogate whether other rRNAs were potential targets of DIMT1. Interestingly, as shown in (**Table 1)**, our RiboMethSeq data show that the methylation of 28S and 5S rRNA was also impaired upon DIMT1-deficiency. Thus, these rRNAs could also be possible targets of *DIMT1* in β-cells. All methylated sites revealed by the RiboMethSeq of scrambled siRNA-treated and *DIMT1* knock down cells are found as a supplementary table **(sTable1)**. The functional implications of methylation of 28S and 5S rRNA by *DIMT1* are yet to be explored.

**Table-1:** Methylated sites present in control samples but not in samples after DIMT1 knockdown. The RiboMethSeq protocol in (ebook ISBN 978-1-4939-6807-7) was applied to generate transcript reads which were aligned to the human rRNA DNA sequences (5S, 5.8S, 18S, 28S) found in the NCBI reference human genome (GRCh38) using Bowtie 2 (PMID: 22388286). The R package RNAmodR.RiboMethSeq (v. 1.2.0) was used for counting of both 5’ and 3’ ends of mapped reads and calculating the Score Mean and RiboMeth‐seq Score (Riboscore) as defined in (PMID: 25417815, PMID: 30539563). A site with a minimum Score Mean of 0.60 and a minimum RiboMeth‐seq score of 0.75 was considered methylated.

| **Gene** | **Methylated Position** | **Strand** | **Methylated Site** | **RBiocore** |
| --- | --- | --- | --- | --- |
| RNA5S9 | 92 | – | Cm | 0,8058252 |
| RNA5S9 | 4 | – | Um | 0,7700288 |
| RNA28SN2 | 2397 | + | Um | 0,7839793 |
| RNA28SN2 | 2403 | + | Um | 0,9242122 |
| RNA28SN2 | 2420 | + | Um | 0,8673416 |
| RNA28SN2 | 2465 | + | Am | 0,7885364 |
| RNA28SN4 | 3333 | + | Cm | 0,8511263 |
| RNA28SN4 | 4978 | + | Um | 0,8689244 |
| RNA5S9 | 35 | - | Um | 0.7962092 |

## Discussion

rRNA modification is highly conserved across species; only a handful of organisms are known to lack it (Ramakrishnan, 2002; Sloan *et al*, 2017). The specific location of the modified residues on the rRNA within the ribosome and its conservation across the species are of great importance and likely play a critical role in translational regulation. rRNA processing, which primarily occurs in the nucleolus, requires various associated proteins that directly facilitate ribosomal assembly and transport of the pre-ribosomal complex from nucleolus to the cytoplasm (Henras *et al*, 2015; Tschochner & Hurt, 2003). A high degree of intricate tethering ensures association of rRNAs to ribosomal proteins (r-proteins), which is regulated by the coordination of substrates and additional factors (Zhou *et al*, 2015). This process is mainly governed by two discrete ribosomal subunits, i.e., 60S and 40S rRNA that associate during translation initiation to form the functional ribosome. The ribosomal subunits and the other components of the translational machinery are typically modified by the addition of methyl groups by methyltransferases. These modifications mainly occur on rRNA, transfer RNA (tRNA), messenger RNA (mRNA), translation factors, and r-proteins. Methylation of these RNA components of the ribosomes is crucial for subunit stabilization, structure stability, translational fidelity and ribosome synthesis and consequently for cellular protein synthesis (Agris, 2015; Roundtree *et al*, 2017).

Here, we report that N^6−^N^6^ dimethylation of two adenosine residues on 18S rRNA by DIMT1 is part of the control of β-cell translation. Interestingly, our data revealed a dependence of methylation executed by DIMT1 on protein synthesis. DIMT1 is highly conserved among different species, which suggests that it very likely has a specific functional role in cells in general, including pancreatic β-cells. We have therefore assessed the role of DIMT1 in β-cells, and demonstrate that methylation of rRNA by DIMT1 is critical for ribosomal subunit assembly. Two of the important proteins associated with 60S and 40S rRNA, (NOB1 and PES1) were found to be altered in DIMT1-deficient cells. Of note, pre-40S ribosome biogenesis in humans requires cleavage by NOB1 at the 3’end of 18S rRNA (Garcia-Gomez *et al*, 2014). This step is of importance to ensure the production of properly assembled and mature 40S rRNA subunits. We found that DIMT1 is essential for β-cell protein translation: loss of *DIMT1* significantly attenuated protein synthesis. The mechanism by which DIMT1 impacts protein synthesis is still not fully resolved but our study suggests that loss of *DIMT1* leads to perturbed ribosomal assembly, in part, owing to a reduced interaction of NOB-1 and PES-1. In yeast, reports have also suggested that regulation of NOB-1 mediates cleavage of site D in the pre-18S rRNA, which is mediated by ITS1 (Internal transcribed spacer). This facilitates Nob-1 access to its cleavage site (Fatica *et al*, 2003; Lamanna & Karbstein, 2009, 2011). The yeast homolog of DIMT1, Dim2, has also been shown to interact with Nob-1(Woolls *et al*, 2011), assisting in rRNA processing. Defects in pre-RNA processing could lead to cell cycle arrest and apoptosis followed by ribosomopathies (Tafforeau *et al*, 2013). Interestingly, our data in human β-cells demonstrated regulation of ribosomal biogenesis by NOB-1 and PES-1 mediated by DIMT1. High-throughput experimental strategies such as mass spectrometric analyses (e.g., SILAC) and ribosome profiling could provide additional information about how these modifications regulate protein synthesis.

Interestingly, the DIMT1 homolog, TFB1M, which is a nuclear-encoded protein, has previously been reported to control protein translation in mitochondria (Sharoyko *et al.*, 2014). This notion further prompted us to characterize the functional implication of DIMT1 in β-cells. Here, we show that knock down of *DIMT1* in rodent and a human β-cell line resulted in impaired insulin secretion and content. Since DIMT1 methylates 18S rRNA in the cytoplasm, we examined whether effects on cytosolic rRNA are translated to mitochondria. To this end, we investigated mitochondrial function; we chose four parameters of mitochondrial function (OXPHOS, mtATP, ΔΨm and mOCR) and assessed whether they were affected by DIMT1-deficiency. We found that deficiency of *DIMT1* in β-cells reduces the levels of mitochondrial-encoded proteins, which most likely is due to attenuation of protein synthesis. This effect could be explained by the observed reduced adenosine dimethylation of the 18S rRNA of the 40S subunit in the cytoplasm in *DIMT1*-silenced cells, which is expected to result in ribosomal subunit destabilization. In addition, there was a significant impact of DIMT1*-*deficiency on both mitochondrial and nuclear-encoded mitochondrial complex subunits. Our data suggest that defects in cytosolic 18S rRNA led to a perturbation of mitochondrial complexes and a nuclear response; together, the cytosolic and mitochondrial impact may lead to the overall decrease in mitochondrial function. It has also been shown that mitochondrial-encoded proteins stabilize some nuclear-encoded proteins (Ali *et al*, 2019). It would be interesting to investigate the interdependency of these two axes, which remains to be fully explored.

In view of these findings, we examined whether defects in mitochondrial complexes translate into downstream effects. As shown in (Figure 3A-B), *DIMT1* knock down affected the complex V (ATP Synthase); we found impaired mitochondrial function illustrated by dissipated ΔΨm, reduced levels of mtATP and reduced mOCR. These impairments in mitochondrial function are most likely due to reduced electron transport and subsequently OXPHOS, leading to mitochondrial dysfunction in *DIMT1*-deficient EndoC-βH1 cells. As a result, β-cell stimulus-secretion coupling was perturbed followed by reduction in insulin release and content. Our findings identify β-cell protein translation as a crucial regulatory process to generate protein required for mitochondrial and cellular function. When impacted, as we observed in DIMT1-deficiency, these processes could lead to impaired protein synthesis, mitochondrial dysfunction and impaired insulin secretion. Our results are in line with our previous observations on the role of TFB1M in β-cells and its involvement in the pathogenesis of T2D (Sharoyko *et al.*, 2014).

Here, we characterized the functional relevance of *DIMT1* methylation as one component of β-cell protein translation. We have shown that DIMT1-deficiency leads to a defect in the ribosomal assembly proteins (NOB1 and PES1) that may lead to attenuation of β-cell protein synthesis. This finding demonstrates the conservation of DIMT1 function in human β-cells, similar to that found in yeast cells by Dim1. In fact, the yeast counterpart of human DIMT1, Dim1, also mediates dimethylation at the 3’-end for pre-ribosomal RNA processing (Lafontaine *et al*, 1995). Dimethylation in other cells, like HeLa, has previously been reported as a critical step in late 18S rRNA modification (Salim & Maden, 1973), but its functional relevance in protein synthesis was not studied. Using primer extension amplifying the flanking region of DIMT1 on 18S rRNA, we detected its dimethylation as a critical process in β-cell function. Several ribosomal assembly factors have been discovered both in yeast and human cells (Tafforeau *et al.*, 2013; Wild *et al*, 2010). The exact role of the 18S rRNA modification in β-cell ribosome function was not completely resolved. To fully understand the diverse and complex pathways of ribosome assembly will require the assessment of the precise function of these methyltransferases, identification of substrates, their binding sites and their functional relevance in cellular pathways. High throughput protein-protein interaction methods, ribosome profiling, and mass spectrometry-based analyses will be helpful to more elucidate β-cell protein translation. Our current working model is shown in (Figure 5). In summary, our work has identified a role of a highly conserved methyltransferase in human β-cell protein translation.

**Figure 5:**
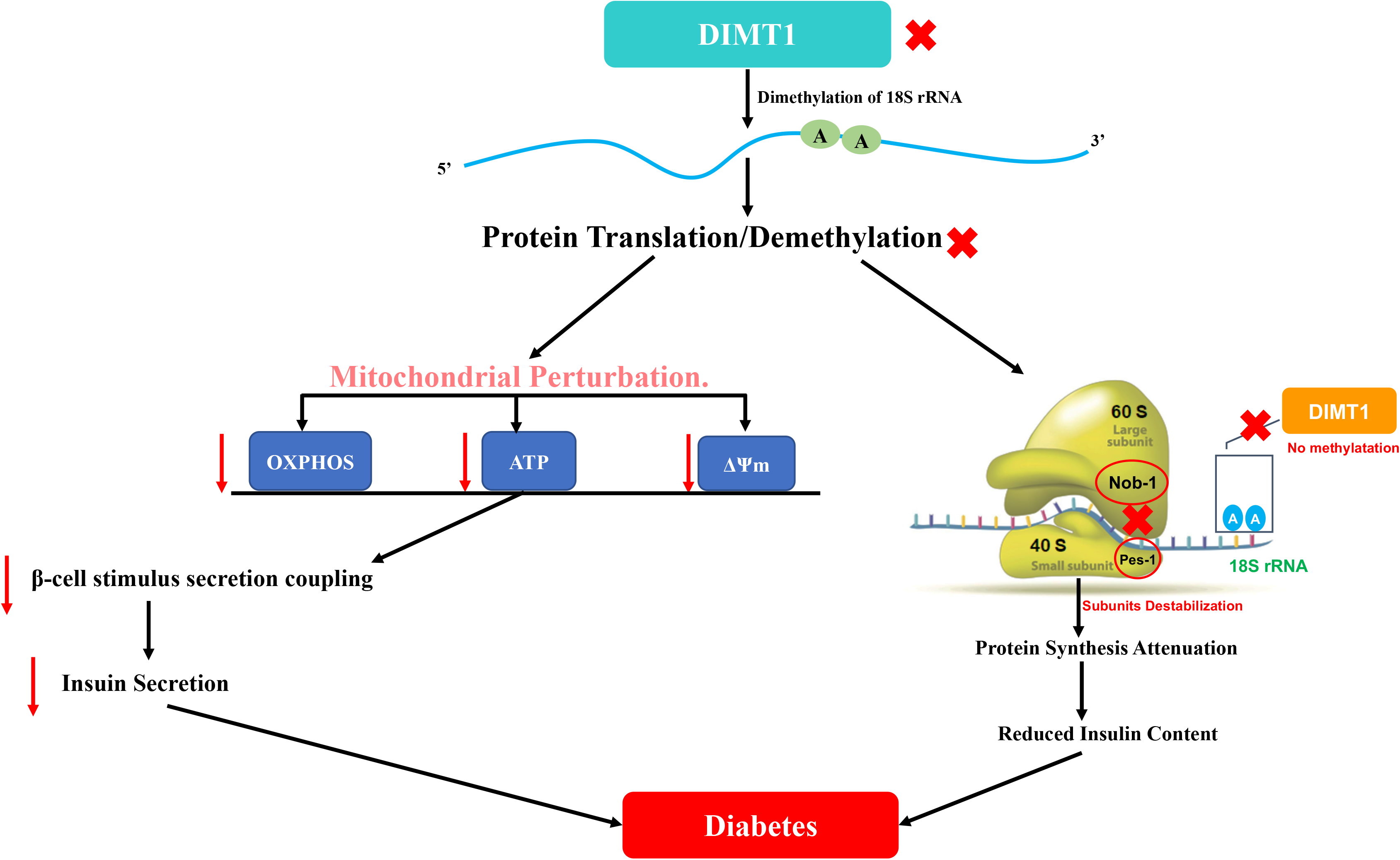
Schematic diagram of DIMT1 mediated β-cell dysfunction. A schematic diagram depicting the mechanisms on how deficiency of *DIMT1* may lead to mitochondrial dysfunction and attenuation of protein synthesis, potentially relevant for diabetes pathophysiology. ATP–Adenosintriphosphate; ΔΨ–Mitochondrial Membrane Potential; OXPHOS – Mitochondrial Oxidative Phosphorylation; NOB-1 – NIN1 (RPN12) Binding Protein 1 Homolog, PES-1 – Pescadillo Ribosomal Biogenesis Factor 1, rRNA – Ribosomal RNA.

## Author contributions

GV designed and performed experiments. Data were also analysed by GV followed by drafting of the manuscript. JE and AK provided the bioinformatics analysis. LRC performed the Seahorse experiments. AB, AH, CL and EC provided feedback on data interpretation. MF contributed to conceptual discussion. HM conceived the project and co-wrote the manuscript. All the authors have reviewed and approved the manuscript.

## Acknowledgements

The authors thank Lund University of Diabetes Centre, Lund University, Malmo, Sweden. The work is supported by European Union’s Horizons 2020 research and innovation program under grant agreement No. 667191. It was also funded by Swedish Research Council (Dnr 2009-1039), LUDC-IRC (Dnr IRC15-0067) and LUDC (349-2006-237). Laila Jacobsson is acknowledged for providing all the laboratory facilities. The authors thank Peter Spégel for input on the proteomics study.

## Conflict of interests

The authors have no competing interests to declare.

## Appendix A. Supplementary data

The following is the supplementary data related to this article:

1. List of the methylated sites in control and knock down cells (S1)
2. List of the differentially altered genes from RNA Sequencing (S2).
3. List of the genes analysed by PANTHER pathways (Supplementary Fig.1)

## Supplementary Figure S1

### PANTHER Pathways database analysis of significantly altered genes

A PANTHER pathway database analysis was performed with a total of 48 significantly expressed genes, which showed significant group differences; this identified 27 molecular functional pathways S1(A) and 55 biological process hits S1(B). The topmost pathways shown emerged from binding, catalytic activity and translation regulator in molecular function S1(A), whereas the maximum hits in biological processes were from biological phase, regulation, cellular biogenesis, localization and metabolic process S1(B). For the full index lists of the genes by Ribomethseq from which the 48 significantly altered genes were selected for Panther pathway categories, see (Supplementary Table S2).

## Notes

### Competing Interest Statement

The authors have declared no competing interest.

## References

Agris PF (2015) The importance of being modified: an unrealized code to RNA structure and function. RNA 21: 552–554

Ali AT, Boehme L, Carbajosa G, Seitan VC, Small KS, Hodgkinson A (2019) Nuclear genetic regulation of the human mitochondrial transcriptome. Elife 8

Aschenbrenner J, Marx A (2016) Direct and site-specific quantification of RNA 2’-O-methylation by PCR with an engineered DNA polymerase. Nucleic Acids Res 44: 3495–3502

Ashcroft FM (2005) ATP-sensitive potassium channelopathies: focus on insulin secretion. J Clin Invest 115: 2047–2058

Basu A, Dalla Man C, Basu R, Toffolo G, Cobelli C, Rizza RA (2009) Effects of type 2 diabetes on insulin secretion, insulin action, glucose effectiveness, and postprandial glucose metabolism. Diabetes care 32: 866–872

Brun T, Maechler P (2016) Beta-cell mitochondrial carriers and the diabetogenic stress response. Biochim Biophys Acta 1863: 2540–2549

Cavaghan MK, Ehrmann DA, Polonsky KS (2000) Interactions between insulin resistance and insulin secretion in the development of glucose intolerance. J Clin Invest 106: 329–333

Cotney J, Shadel GS (2006) Evidence for an early gene duplication event in the evolution of the mitochondrial transcription factor B family and maintenance of rRNA methyltransferase activity in human mtTFB1 and mtTFB2. J Mol Evol 63: 707–717

Fatica A, Oeffinger M, Dlakic M, Tollervey D (2003) Nob1p is required for cleavage of the 3’ end of 18S rRNA. Mol Cell Biol 23: 1798–1807

Fex M, Nicholas L.M, Vishnu N, Medina A, Sharoyko V.V, Nicholls D.G, Spégel P, Mulder H (2018) The pathogenetic role of β-cell mitochondria in type 2 diabetes.J of Endocrinology 236: R145–R159

Gandasi NR, Yin P, Riz M, Chibalina MV, Cortese G, Lund PE, Matveev V, Rorsman P, Sherman A, Pedersen MG et al (2017) Ca^2+^ channel clustering with insulin-containing granules is disturbed in type 2 diabetes. J Clin Invest 127: 2353–2364

Garcia-Gomez JJ, Fernandez-Pevida A, Lebaron S, Rosado IV, Tollervey D, Kressler D, de la Cruz J (2014) Final pre-40S maturation depends on the functional integrity of the 60S subunit ribosomal protein L3. PLoS Genet 10: e1004205

Guja KE, Venkataraman K, Yakubovskaya E, Shi H, Mejia E, Hambardjieva E, Karzai AW, Garcia-Diaz M (2013) Structural basis for S-adenosylmethionine binding and methyltransferase activity by mitochondrial transcription factor B1. Nucleic Acids Res 41: 7947–7959

Henras AK, Plisson-Chastang C, O’Donohue MF, Chakraborty A, Gleizes PE (2015) An overview of pre-ribosomal RNA processing in eukaryotes. Wiley Interdiscip Rev RNA 6: 225–242

Hohmeier HE, Mulder H, Chen G, Henkel-Rieger R, Prentki M, Newgard CB (2000) Isolation of INS-1-derived cell lines with robust ATP-sensitive K+ channel-dependent and -independent glucose-stimulated insulin secretion. Diabetes 49: 424–430

Koeck T, Olsson AH, Nitert MD, Sharoyko VV, Ladenvall C, Kotova O, Reiling E, Ronn T, Parikh H, Taneera J et al (2011) A common variant in TFB1M is associated with reduced insulin secretion and increased future risk of type 2 diabetes. Cell metabolism 13: 80–91

Lafontaine D, Vandenhaute J, Tollervey D (1995) The 18S rRNA dimethylase Dim1p is required for pre-ribosomal RNA processing in yeast. Genes Dev 9: 2470–2481

Lamanna AC, Karbstein K (2009) Nob1 binds the single-stranded cleavage site D at the 3’-end of 18S rRNA with its PIN domain. Proc Natl Acad Sci U S A 106: 14259–14264

Lamanna AC, Karbstein K (2011) An RNA conformational switch regulates pre-18S rRNA cleavage. J Mol Biol 405: 3–17

Lee Y, Fluckey JD, Chakraborty S, Muthuchamy M (2017) Hyperglycemia- and hyperinsulinemia-induced insulin resistance causes alterations in cellular bioenergetics and activation of inflammatory signaling in lymphatic muscle. FASEB J 31: 2744–2759

Maechler P, Li N, Casimir M, Vetterli L, Frigerio F, Brun T (2010) Role of mitochondria in beta-cell function and dysfunction. Advances in experimental medicine and biology 654: 193–216

Metodiev MD, Lesko N, Park CB, Camara Y, Shi Y, Wibom R, Hultenby K, Gustafsson CM, Larsson NG (2009) Methylation of 12S rRNA is necessary for in vivo stability of the small subunit of the mammalian mitochondrial ribosome. Cell metabolism 9: 386–397

Nicholas LM, Valtat B, Medina A, Andersson L, Abels M, Mollet IG, Jain D, Eliasson L, Wierup N, Fex M et al (2017) Mitochondrial transcription factor B2 is essential for mitochondrial and cellular function in pancreatic beta-cells. Mol Metab 6: 651–663

Ramakrishnan V (2002) Ribosome structure and the mechanism of translation. Cell 108: 557–572

Rorsman P, Braun M, Zhang Q (2012) Regulation of calcium in pancreatic alpha- and beta-cells in health and disease. Cell Calcium 51: 300–308

Roundtree IA, Evans ME, Pan T, He C (2017) Dynamic RNA Modifications in Gene Expression Regulation. Cell 169: 1187–1200

Salim M, Maden BE (1973) Early and late methylations in HeLa cell ribosome maturation. Nature 244: 334–336

Schuit F, De Vos A, Farfari S, Moens K, Pipeleers D, Brun T, Prentki M (1997) Metabolic fate of glucose in purified islet cells. Glucose-regulated anaplerosis in beta cells. The Journal of biological chemistry 272: 18572–18579

Seidel-Rogol BL, McCulloch V, Shadel GS (2003) Human mitochondrial transcription factor B1 methylates ribosomal RNA at a conserved stem-loop. Nat Genet 33: 23–24

Shanik MH, Xu Y, Skrha J, Dankner R, Zick Y, Roth J (2008) Insulin resistance and hyperinsulinemia: is hyperinsulinemia the cart or the horse? Diabetes care 31 Suppl 2: S262–268

Sharoyko VV, Abels M, Sun J, Nicholas LM, Mollet IG, Stamenkovic JA, Gohring I, Malmgren S, Storm P, Fadista J et al (2014) Loss of TFB1M results in mitochondrial dysfunction that leads to impaired insulin secretion and diabetes. Hum Mol Genet 23: 5733–5749

Sloan KE, Warda AS, Sharma S, Entian KD, Lafontaine DLJ, Bohnsack MT (2017) Tuning the ribosome: The influence of rRNA modification on eukaryotic ribosome biogenesis and function. RNA Biol 14: 1138–1152

Tafforeau L, Zorbas C, Langhendries JL, Mullineux ST, Stamatopoulou V, Mullier R, Wacheul L, Lafontaine DL (2013) The complexity of human ribosome biogenesis revealed by systematic nucleolar screening of Pre-rRNA processing factors. Mol Cell 51: 539–551

Tschochner H, Hurt E (2003) Pre-ribosomes on the road from the nucleolus to the cytoplasm. Trends Cell Biol 13: 255–263

Wild T, Horvath P, Wyler E, Widmann B, Badertscher L, Zemp I, Kozak K, Csucs G, Lund E, Kutay U (2010) A protein inventory of human ribosome biogenesis reveals an essential function of exportin 5 in 60S subunit export. PLoS Biol 8: e1000522

Wollheim CB, Maechler P (2002) Beta-cell mitochondria and insulin secretion: messenger role of nucleotides and metabolites. Diabetes 51 Suppl 1: S37–42

Woolls HA, Lamanna AC, Karbstein K (2011) Roles of Dim2 in ribosome assembly. The Journal of biological chemistry 286: 2578–2586

Zhou X, Liao WJ, Liao JM, Liao P, Lu H (2015) Ribosomal proteins: functions beyond the ribosome. J Mol Cell Biol 7: 92–104

Zorbas C, Nicolas E, Wacheul L, Huvelle E, Heurgue-Hamard V, Lafontaine DL (2015) The human 18S rRNA base methyltransferases DIMT1L and WBSCR22-TRMT112 but not rRNA modification are required for ribosome biogenesis. Mol Biol Cell 26: 2080–2095

