## Supplementary figures and images for "DIMT1, a regulator of ribosomal biogenesis, controls β-cell protein synthesis, mitochondrial function and insulin secretion"

### Supplementary Figure 1

#### Slide 1
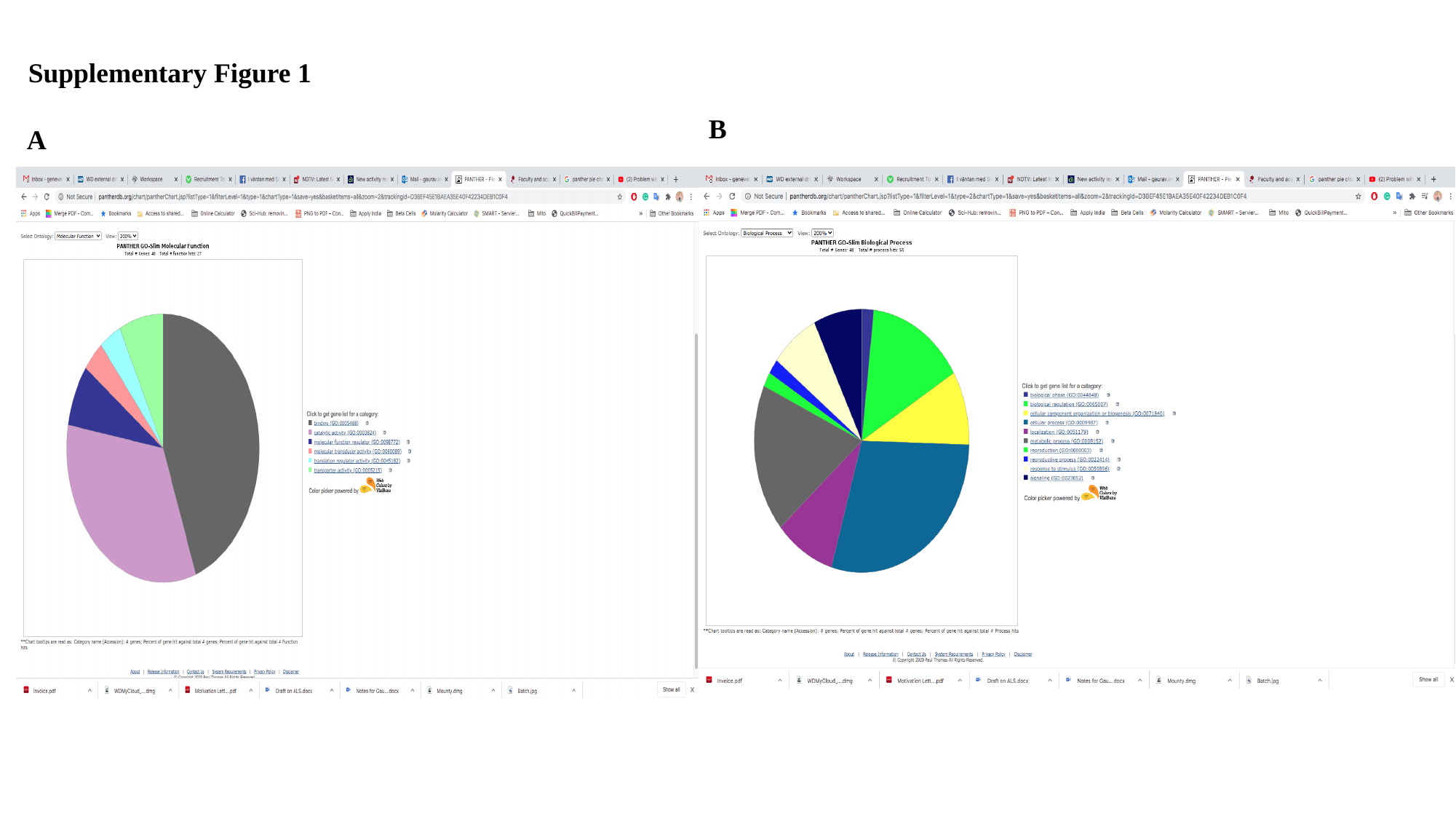

Supplementary Figure 1
B
A
